# Global biogeography of *Tetragnatha* spiders reveals multiple colonization of the Caribbean

**DOI:** 10.1101/452227

**Authors:** Klemen Čandek, Ingi Agnarsson, Greta J. Binford, Matjaž Kuntner

## Abstract

Organismal variation in dispersal ability can directly affect levels of gene flow amongst populations, therefore importantly shaping species distributions and species richness patterns. The intermediate dispersal model of biogeography (IDM) predicts that in island systems, species diversity of those lineages with an intermediate dispersal potential is the highest. We broadly test this prediction, focusing on ‘four-jawed spiders’ (genus *Tetragnatha*) of the Caribbean archipelago. First, we report on original sampling of this globally distributed genus with numerous widespread as well as endemic species. We then reconstruct multiple *Tetragnatha* phylogenies from roughly 300 individuals delineated into 54 putative species. Our results support the monophyly of the four-jawed spiders but reject the monophyly of those lineages that reach the Caribbean, where we find low levels of endemism yet high diversity within *Tetragnatha*. The chronogram detects a potential early overwater colonization of the Caribbean, and in combination with reconstructed biogeographic history, refutes the possibility of ancient vicariant origins of Caribbean *Tetragnatha* as well as the GAARlandia land-bridge scenario. Instead, biogeographic results hypothesize multiple colonization events to, and from the Caribbean since mid-Eocene to late-Miocene. *Tetragnatha* seems unique among the arachnids explored so far in comprising some species that are excellent dispersers, and others that are not, perhaps having secondarily lost this dispersal propensity. A direct test of the IDM would require consideration of three categories of dispersers. However, four-jawed spiders do not fit one of these three a *priori* definitions, but rather represent a more complex combination of attributes of a ‘dynamic disperser’.

## 1 Introduction

Species distributions and species richness can vastly vary among taxonomic units of comparable ranks. Evolutionary biology has long been trying to understand which factors contribute to such variation (Kozak and Wiens, 2016a, 2016b). On one hand, abiotic factors such as habitable area size, climate conditions or the presence of barriers may contribute to this variation (Rabosky and Hurlbert, 2015; Schluter and Pennell, 2017). On the other hand, biological attributes such as species generation time (Rajakaruna and Lewis, 2018), clade age (Marin and Hedges, 2016) and species dispersal ability (Borda-de-Água et al., 2017; Lenoir et al., 2012) may be equally important. Dispersal ability, in particular, has the potential to directly affect levels of gene flow amongst populations and species’ potential to reach new habitats.

A low dispersal potential can limit colonization success and gene flow amongst populations, while high dispersal potential enables colonization of remote areas and maintains higher levels of gene flow. In theory, both of these extreme cases (low and high dispersal) constrain the number of speciation events (Agnarsson et al., 2014; Claramunt et al., 2012). The intermediate dispersal model (Agnarsson et al., 2014; Claramunt et al., 2012) predicts that in island systems, the species diversity of those lineages with an intermediate dispersal potential is the highest.

Taxa that contain species with poor dispersal abilities are associated with high levels of endemism, which is often associated with biogeographic patterns of multiple single island endemics across archipelagos (Opatova and Arnedo, 2014; Pitta et al., 2014). On the other hand, taxa that contain species with excellent dispersal potential are usually widely, sometimes globally, distributed (Essl et al., 2013; Kuntner and Agnarsson, 2011). Studying and understanding biogeography of species and higher taxonomic ranks on local as well as on global scales could provide insights to a clade’s intrinsic propensities to disperse. Island systems have long been a model for such biogeographic research on a local scale (Darwin, 1859; Warren et al., 2015). Islands are discrete units with different degrees of barriers to gene flow for terrestrial organisms leading to different degrees of adaptation and diversification (Reynolds et al., 2016; Ricklefs and Bermingham, 2008). Extreme forms of morphological adaptations or even secondary loss of dispersal abilities in putatively good dispersers are not unusual in island systems (Frankham, 2008; Gillespie et al., 1997; Meiri, 2017). While some species undergo rapid local adaptation and diversification, others readily disperse among islands and form a wide distribution across many islands of the archipelago and possibly beyond. However, these widely distributed species, are sometimes found to be species complexes rather than a single species (Cornils et al., 2017; Tomioka et al., 2016). Therefore, a close examination of the biogeographic pattern and population structure of such taxa can help with the discovery of new or cryptic species.

Spiders are becoming popular in biogeographic research (Agnarsson et al., 2018, 2016; Chamberland et al., 2018; Cotoras et al., 2018; Gillespie et al., 2018), and are particularly suitable as models due to the breath of their taxonomic, genetic, evolutionary, and biological diversity. Spiders represent an ancient lineage that today comprises over 47,000 described, and likely many more, species from 117 families (WSC, 2018) inhabiting most terrestrial ecosystems. Interestingly, various taxonomic groups of comparable ranks within spiders exhibit highly variable dispersal abilities, degrees of endemism, species richness and species distribution (Foelix, 2011).

Our study focuses on ‘four-jawed spiders’ (genus *Tetragnatha*, family Tetragnathidae), with special emphasis on the Caribbean archipelago. This diverse genus includes 349 species and has been extensively used in biogeographic research, notably in Hawaii and other Pacific archipelagos (Casquet et al., 2015; Cotoras et al., 2017; Gillespie, 2005, 2003). While some species are single island endemics, others show extremely wide, even cosmopolitan distributions e.g. *T. nitens* (WSC, 2018). *Tetragnatha* is generally considered to have an extraordinary dispersal potential and is able to quickly reach even the most remote, newly formed islands (Cotoras et al., 2017). This assumption, reinforced by a study of Okuma, C. and Kisimoto, R. (1981), found that 96 % of the aerial plankton collected 400 km off the Chinese shore were *Tetragnatha* spiders, passively dispersing with behavior called ballooning (Bell et al., 2005; Foelix, 2011).

We studied a rich original collection of *Tetragnatha* from the Caribbean archipelago. With species delimitation method we estimate to have 25 putative species in our original dataset. We then reconstruct the evolutionary history of four-jawed spiders of the Caribbean using a mitochondrial COI gene and a nuclear 28S gene fragment. We place the Caribbean phylogeny into a global context by adding a single sequence for every available published *Tetragnatha* species on GenBank, increasing the number of putative species in Bayesian analysis to 54. Moreover, we estimate the number of, the timing, and the directionality of all Caribbean colonization events by *Tetragnatha*, then look for potential agreement with common biogeographic scenarios on the Caribbean e.g. colonization by overwater dispersal or vicariant origins (Hedges, 2006) or a combined scenario, GAARlandia (Ali, 2012; Iturralde-Vinent and MacPhee, 1999). We contrast the biogeography of *Tetragnatha* with those of other spider lineages on the Caribbean, including a close relative *Cyrtognatha*, and broadly estimate dispersal abilities of *Tetragnatha* in the context of intermediate dispersal model (IDM). Finally, we examine a population structure of a well-represented “species” from the Caribbean by reconstructing its haplotype network and explore the potential of multiple cryptic species, rather than a single widely distributed species, comprising this taxon.

## 2 Material and Methods

### 2.1 Material acquisition

We collected the material for our research as a part of large-scale Caribbean biogeography (CarBio) project. We used standard methods for collecting spiders (Agnarsson et al., 2013; Coddington et al., 1991), namely day- and night-time beating and visual aerial search. We fixed the collected material in 96 % ethanol at the site of field work and stored it at −20/-80 °C. We then used light microscopy to verify the genus and to identify species, where possible.

### 2.2 Molecular procedures

We isolated DNA using QIAGEN DNeasy Tissue Kit (Qiagen, Inc., Valencia, CA) at the University of Vermont (Vermont, USA), or an Autogenprep965 automated phenol chloroform extraction at the Smithsonian Institution (Washington, DC, USA), or a robotic DNA extraction with Mag MAX™ Express magnetic particle processor Type 700 with DNA Multisample kit (Applied Biosystems, Foster City, CA, USA), following modified protocols (Vidergar et al., 2014) at the EZ Lab (Ljubljana, Slovenia).

We targeting two genetic markers, a mitochondrial (COI) and a nuclear one (28S rRNA). We used the forward LCO1490 (GGTCAACAAATCATAAAGATATTGG) and the reverse C1-N-2776 (GGATAATCAGAATATCGTCGAGG) for COI amplification. The 25 µL reaction volume contained the mixture of: 5 µL Promega‘s GoTaq Flexi Buffer, 0.15 µL GoTaq Flexi Polymerase, 0.5 µL dNTP’s (2 mM each, Biotools), 2.3 µL MgCl2 (25 mM, Promega), 0.5 µL of each primer (20 µM), 0.15 µL BSA (10 mg/mL; Promega), 2 µL DNA template and sterile distilled water for the remaining volume. We set PCR cycling protocol as follows: initial denaturation (5 min at 94 °C), 20 repeats (60 s at 94°C, 90 s at 44 °C while increasing 0.5 ° per repeat, 1 min at 72 °C), 15 repeats (90 s at 94 °C, 90 s at 53.5 °C, 60 s at 72 °C), final elongation (7 min at 72 °C).

We used the forward 28Sa (also known as 28S-D3A; GACCCGTCTTGAAACACGGA) and the reverse 28S-rD5b (CCACAGCGCCAGTTCTGCTTAC) for 28S amplification. The 35 µL reaction volume contained the mixture of: 7.1 µL Promega‘s GoTaq Flexi Buffer, 0.2 µL GoTaq Flexi Polymerase, 2.9 µL dNTP’s (2 mM each, Biotools), 3.2 µL MgCl2 (25 mM, Promega), 0.7 µL of each primer (20 µM), 0.2 µL BSA (10 mg/mL; Promega), 1 µL DNA template and sterile distilled water for the remaining volume. We set PCR cycling protocol as follows: initial denaturation (7 min at 94 °C), 20 repeats (45 s at 96 °C, 45 s at 62 °C while decreasing 0.5 ° per repeat, 1 min at 72 °C), 15 repeats (45 s at 96 °C, 45 s at 52 °C, 60 s at 72 °C), final elongation (10 min at 72 °C).

We used Geneious v. 5.6.7 (Kearse et al., 2012) for de-novo sequence assembly and initial manipulation. We used a combination of MEGA (Tamura et al., 2013) and Mesquite (Maddison and Maddison, 2008) for basic sequence analysis, renaming and concatenating matrices of both genetic markers. We then used the online version of MAFFT (Katoh and Standley, 2013) for sequence alignment.

We obtained 254 original COI and 54 original 28S sequences of *Tetragnatha* spiders. We incorporated an additional 45 *Tetragnatha* COI sequences from GenBank, representing all published sequences (1 per species, except 3 × *T. nitens*, 2 × *T. shoshone*, 2 × *T. viridis*, 2 × *T. maxillosa*) of sufficient quality (over 70 % of overlap with our sequences). For outgroups, we used nine COI and seven 28S originally generated sequences from non-*Tetragnatha* genera as well as ten COI and five 28S sequences mined from GenBank. Altogether, our broadest dataset comprised 318 COI sequences and 66 28S sequences. Our specimen selection focused on the Caribbean but our broad taxon sampling ensured a global representation of *Tetragnatha*. The concatenated matrix for COI and 28S genes is 1199 nucleotide long, 649 for COI and 550 for 28S. Relevant specimen details and GenBank accession codes are presented in Table A.1.

### 2.3 Species delimitation

We used Automatic Barcode Gap Discovery (ABGD) (Puillandre et al., 2012) for estimating the number of molecular operational taxonomic units (MOTUs) using all COI sequence in the dataset. We set Pmin to 0.001 and Pmax to 0.2 with 30 steps between those values. We set the X (relative gap width) to different values from 1.5 to 3 to check for the consistency of the results.

### 2.4 Phylogenetic analyses

#### 2.4.1 Two gene, species level phylogeny

We used a subset of our data based on the results from the above described species delimitation analysis to create a concatenated matrix of two gene markers (COI and 28S) for Bayesian phylogenetic reconstruction. We used 54 of the *Tetragnatha* individuals collected for this work that represent 20 MOTUs and added 12 sequences as outgroups (Table A.1). We used MrBayes (Huelsenbeck and Ronquist, 2001) to run two independent runs, each with four MCMC chains, for 30 million generations. We partitioned the dataset per genetic marker, set a sampling frequency of 2000 and set a relative burn-in to 25 %. We used Generalised time-reversible model with gamma distribution and invariant sites (GTR+G+I) as a model of nucleotide substitution for both partitions, as suggested by AIC and BIC criterion from preceding analyses of our sequences with jModelTest2 (Darriba et al., 2012). The starting tree was random.

We examined the statistical parameters and MCMC chains convergence with sump command within MrBayes and with Tracer (Rambaut et al., 2018). Trees were visualized with FigTree (Bouckaert et al., 2014).

#### 2.4.1 All-terminal, single gene phylogeny

Using MrBayes, we reconstructed two Bayesian phylogenies using all available *Tetragnatha* terminals - all originally sampled *Tetragnatha* (N = 254) and a single COI sequence per every available, published, *Tetragnatha* species (N = 45). We added 19 specimens as outgroups (Table A.1). We included multiple Hawaiian *Tetragnatha* species belonging to “spiny-leg” clade, to serve as a check, since their monophyly is well established (Casquet et al., 2015; Gillespie et al., 1997) and should thus be recovered in our own phylogenetic reconstructions.

The first all-terminal phylogenetic reconstruction used an unconstrained approach while we enforced the *Tetragnatha* monophyly for the second one. The settings for both analyses were as in the above species level analysis although the number of MCMC generations was increased to 100 million. Additionally, we set the parameter “contype” within the “sumt” command to “allcompat” to get a fully resolved tree regardless of potential low support for some internal nodes.

We then tested which model, the unconstrained or the constrained, better fits our data by comparing marginal likelihood scores and by Tracer model comparison analysis. We examined the statistical parameters and MCMC chains convergence with sump command within MrBayes and with Tracer. Trees were visualized with FigTree.

#### 2.4.2 Caribbean Tetragnatha monophyly testing

To test whether *Tetragnatha* from the Caribbean are monophyletic, we ran two additional Bayesian analyses. We created a subset of COI sequences, a single for each MOTU/recognized species. We then ran the unconstrained analysis and compared it to the analysis with enforced monophyly of Caribbean endemic *Tetragnatha* species. To confirm or reject the monophyly of *Tetragnatha* species present exclusively on the Caribbean islands, we compared the marginal likelihood scores between the models and ran a Tracer model comparison analysis. The general settings of both (constrained and unconstrained) phylogenetic analyses were as in the above described species level analysis although the number of MCMC generations were increased to 50 million.

#### 2.4.3 Time-calibrated phylogenetic reconstruction

We used BEAST (Bouckaert et al., 2014) for a time-calibrated phylogeny. We created a subset of COI sequences as per BEAST requirements: a single for each MOTU/recognized species instead of multiple terminals per species with zero or low divergence between them (Table A.1). We used bModelTest (Bouckaert and Drummond, 2017) expansion in BEAST as a substitution model in model selection tab. Considering model comparison results, we followed Bidegaray-Batista & Arnedo (2011). We selected a relaxed log-normal molecular clock and set the priors accordingly: ucld.mean with normal distribution, mean value of 0.0112 and standard deviation 0.001; ucld.stdev with exponential distribution and the mean value of 0.666. We set the number of MCMC generations to 30 million with sampling frequency of 1000. We constrained the whole topology based on the results from the all-terminal COI phylogeny and let BEAST to only estimate branch lengths and associated timing. Furthermore, we added three calibration points to the chronogram. The first was the well supported appearance of the Hawaiian islands at 5.1 MYA, therefore the diversification of Hawaiian clade was constrained with a uniform prior with upper bound at 5.1 MYA and lower bound at 0. The second calibration point is the time of the appearance of Lesser Antillean islands, which are presumably not older than 11 million years (Peck, 2011). Therefore, the diversification of the clade uniting two reasonable Lesser Antillean endemic species (SP5 and SP9) was constrained using a uniform prior with upper bound at 11 MYA and lower bound at 0. The last calibration point originates from spider phylogenomic literature where Garrison et al. (2016) and Bond et al. (2014) estimated the Tetragnathidae age at 100 (44 – 160) and 99 (64 – 133) MYA. Therefore, we set the MRCA prior on the node *Tetragnatha + Arkys* (the latter outgroup in Arkyidae) (Dimitrov et al., 2017) with the following settings: exponential distribution, mean value of 31.5 and 44 offset, corresponding to the soft upper bound at 160 MYA and hard lower bound at 44.8 MYA.

To determine burn-in, check the MCMC convergence and other statistical parameters we used Tracer. We used TreeAnnotator (Bouckaert et al., 2014) to summarize trees with 10% burn-in and node heights set as median heights. We used FigTree for consensus tree visualization.

### 2.5 Ancestral area estimation and biogeographic stochastic mapping

We used BioGeoBEARS (Matzke, 2013) package in R version 3.5.0 (R Core Team, 2018) for biogeographic analyses of *Tetragnatha* from the Caribbean and wider. We used the ultrametric tree from BEAST with outgroups removed. We conducted the analysis with the remaining 54 species/MOTUs and created a geographic data list with two possible areas: The Caribbean (C) and “Other” (O), meaning four possible ancestral states at each node (C, O, CO and ‘null’). We thereby achieved a good resolution for estimating biogeographic events between the Caribbean and non-Caribbean areas. We tested all six possible biogeographical models implemented in BioGeoBEARS: DEC (+J), DIVALIKE (+J) and BAYAREALIKE (+J), testing their suitability for our data with Akaike information criterion (AIC) and sample-size corrected AIC (AICc).

To correctly estimate the number and types of biogeographic events in the Caribbean, we performed biogeographical stochastic mapping (BSM), analysis expansion in BioGeoBEARS. We used the most suitable (BAYAREALIKE+J) model, as discovered in the results from previous analysis. We simulated 100 exact biogeographic histories and extracted estimated number and types of events and presented them as histograms.

### 2.6 *Tetragnatha* SP2 haplotype network

We used pegas (Paradis, 2010) package in R v. 3.5.0 to calculate a haplotype network for *Tetragnatha* SP2, the species selected based on its broad Caribbean distribution and potential for population structuring according to the all-terminal phylogeny. We used 76 individuals and trimmed the sequences to equal lengths of 570 nucleotide each with 100% of overlap, as required by the software. Two sequences of *Tetragnatha* SP2 were discarded due to their short length. The circle sizes within a haplotype network correspond to the frequency of identical haplotypes while the colors represent different geographical areas.

## 3 Results

### 3.1 Molecular procedures and MOTU estimation

We collected 254 individuals from Cuba, Jamaica, Lesser Antilles, Hispaniola, Puerto Rico, SouthEast USA and Central America (Table A.1). We confirmed that all individuals were morphologically *Tetragnatha* and identified eight described species. Additionally, we estimated another 17 MOTUs (labeled SP and number), using ABGD computational method (Note A.1). ABGD was consistent in estimating the number of MOTUs for all tested X (relative gap width) values. Moreover, ABGD detected a 3.8 % wide barcoding gap (measured in K2P (Kimura, 1980) percent distance), closely matching that of a barcoding gap width estimated for Tetragnathidae by Čandek & Kuntner (2014).

ABGD species delimitation correctly separated all species from GenBank with the exception of *T. praedonia* and *T. nigrita*. Those two species are clustered together with only 1.1 % sequence divergence between them, suggesting that one of them is misidentified. Similarly, *T. nitens* is separated into two species, one of them clustering with *T. moua*. This suggests that *T. nitens* is either a species complex, or that it may represent another case of misidentification. Altogether, our originally collected material comprises 25 putative species while together with GenBank sequences our dataset contains 54 putative species. Of those, 11 are Caribbean endemics while 15 species are found both in the Caribbean as well as elsewhere. In our dataset, 28 species are not found in the Caribbean.

### 3.2 Molecular phylogeny

Our two gene, species level, phylogeny (Fig. 1) supports *Tetragnatha* monophyly. A well-supported basal *Tetragnatha* node contains some, but not all *T. shoshone* exemplars, refuting the validity of that MOTU. The remaining species/MOTUs generally group together with high support. Except for a single node, internal nodes in this phylogeny are resolved (Fig. 1).

**Figure 1.**
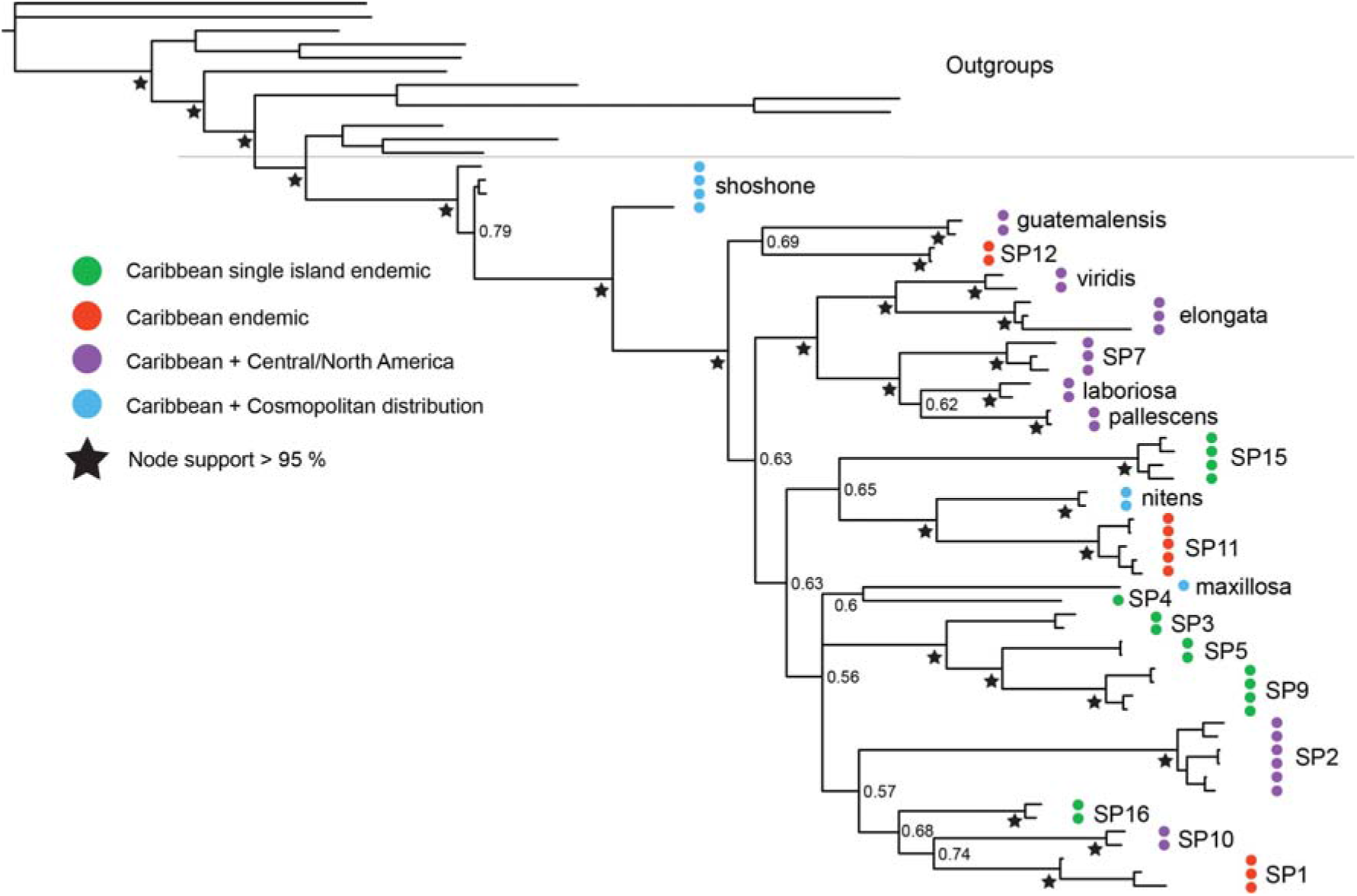
Species level Bayesian phylogeny of the Caribbean *Tetragnatha* based on COI and 28S. Relationships agree with *Tetragnatha* monophyly.

The COI phylogenetic reconstruction in Figure 2 represents the relationships of all originally collected and all data-mined *Tetragnatha*. Because the unconstrained all-terminal phylogeny (Figure A.1) recovers a paraphyletic *Tetragnatha* that also includes *Pachygnatha*, we constrained the *Tetragnatha* monophyly (Fig. 2). Occasional inaccuracies at deeper levels of mitochondrial phylogenies are to be expected due to the high levels of information saturation in COI gene (Brandley et al., 2011). Our approach to constrain the all-terminal COI phylogeny is justified by marginal likelihood scores test with Tracer model comparison (logBF = 710.4 ± 0.061 from 500 bootstrap replicates favoring the constrained model), and moreover, the constrained phylogeny exhibits higher posterior node probabilities. The Hawaiian *Tetragnatha* always formed a well-supported clade, a result giving credibility to our analyses.

**Figure 2.**
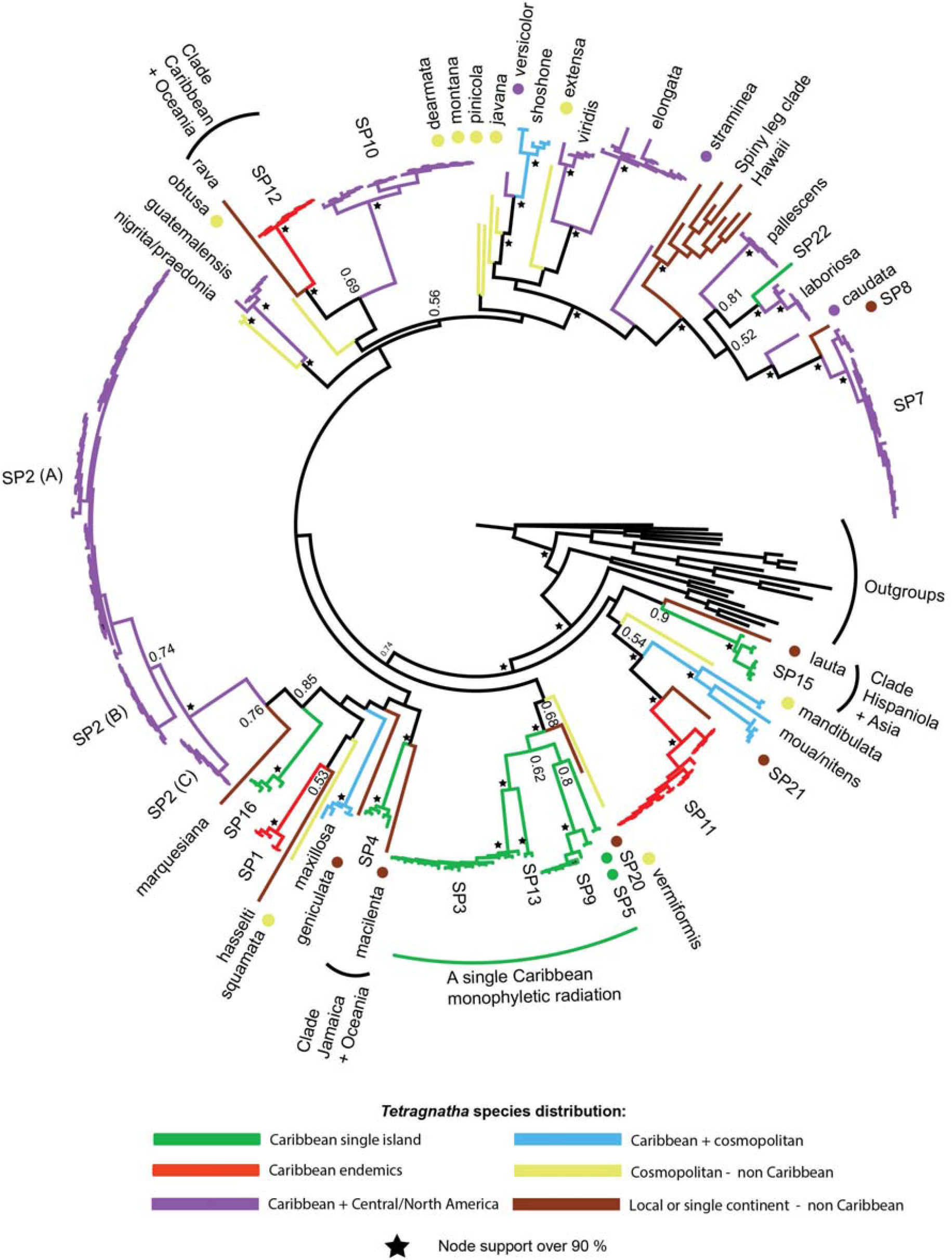
The all-terminal mitochondrial Bayesian phylogeny of the Caribbean and global *Tetragnatha* representatives. The scattered phylogenetic pattern of single island endemic, Caribbean endemic and cosmopolitan species, as well as highly supported clades of geographically distant relatives reveal a profile of a “dynamic disperser”. Note the non-monophyly of Caribbean *Tetragnatha*.

This phylogenetic pattern (Fig. 2) reveals eight MOTUs as Caribbean single island endemics and an additional three that are Caribbean endemics. The remaining putative species have more widespread distributions. Although the phylogenetic pattern in itself already hints at Caribbean *Tetragnatha* not being monophyletic, its monophyly is formally rejected by the marginal likelihood scores testing with Tracer model analysis (logBF = 163.6 ± 0.039 SE from 500 bootstrap replicates favoring unconstrained model). The only multispecies monophyletic Caribbean *Tetragnatha* clade appears to be the group uniting SP3, 5, 9, 13. Because *Tetragnatha* SP2, the species with the highest number of individuals (N = 78) with a distribution across all major Caribbean islands, shows a marked population structuring (labeled as SP2 A-C in Fig. 2), it was selected for subsequent haplotype network analyses.

The examination of p files in Tracer and sump command within MrBayes for all of the above described phylogenies, found high ESS values, the lowest being 12,487 for two gene and 617 for COI only phylogeny. MCMC chain successfully converged for all parameters.

### 3.3 Time calibrated phylogeny

The BEAST chronogram suggests tetragnathids diverged from their sister family Arkyidae (Dimitrov et al., 2017) 58 MYA (million years ago) (95% HPD 44.5 – 81.9) (Fig. 3). We estimate that the diversification of those *Tetragnatha* species that are represented in our dataset began 46 MYA (44 – 53.3). The only multispecies Caribbean endemic clade appeared to have diverged from its sister species from Mexico 20 MYA (12.8 – 29 MYA). Time estimates of the species divergences generally fit the estimated geologic ages of the Caribbean islands.

**Figure 3.**
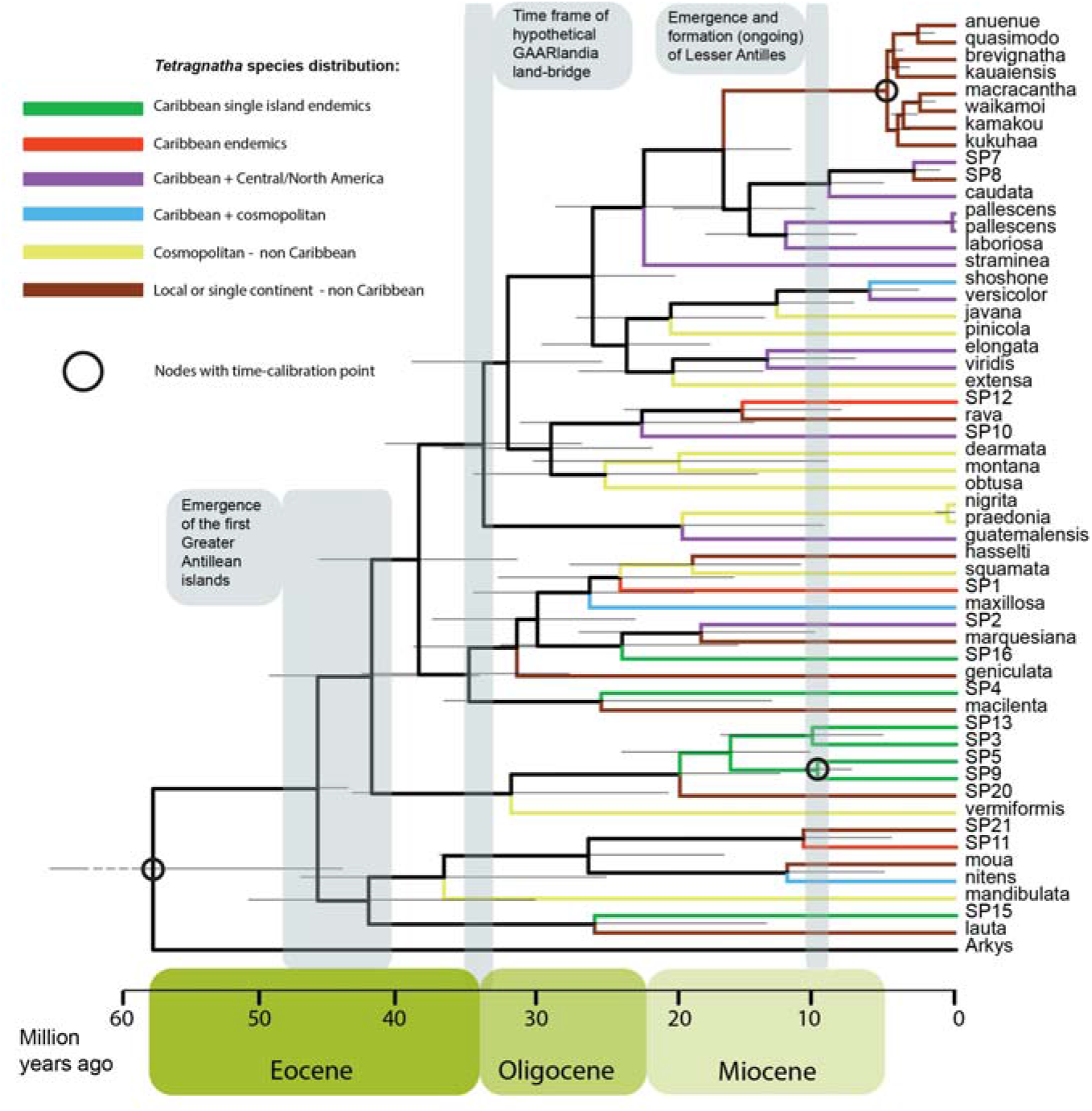
Time-calibrated BEAST phylogeny of *Tetragnatha*. This chronogram allows for an early Caribbean colonization by *Tetragnatha*. The reconstructed timing and pattern favor multiple overwater dispersal events over the GAARlandia or ancient vicariance scenarios. The bars on nodes represent 95 % HPD intervals.

The BEAST phylogeny is well supported as seen from the examination of log files (all EES > 266). Additional examination of log files with bModelAnalyzer revealed the most used nucleotide substitution model was a version of TVM, followed by versions of TN93 and GTR (Fig. A.2) (for details on bModelTest check Bouckaert & Drummond (2017)).

### 3.4 Biogeographic analyses

BioGeoBEARS model comparison recovered the BAYAREALIKE+J model as the most suitable for our data: Model comparison recovered the highest LnL scores for BAYAREALIKE+J, regardless of the scoring criterion (Table A.2). Moreover, it revealed that a founder effect within the model (+J parameter) is significantly better suited for our data than the model without this parameter (p < 0.01). Ancestral area estimation (Fig. 4) hints at multiple and independent origins of the Caribbean *Tetragnatha*.

**Figure 4.**
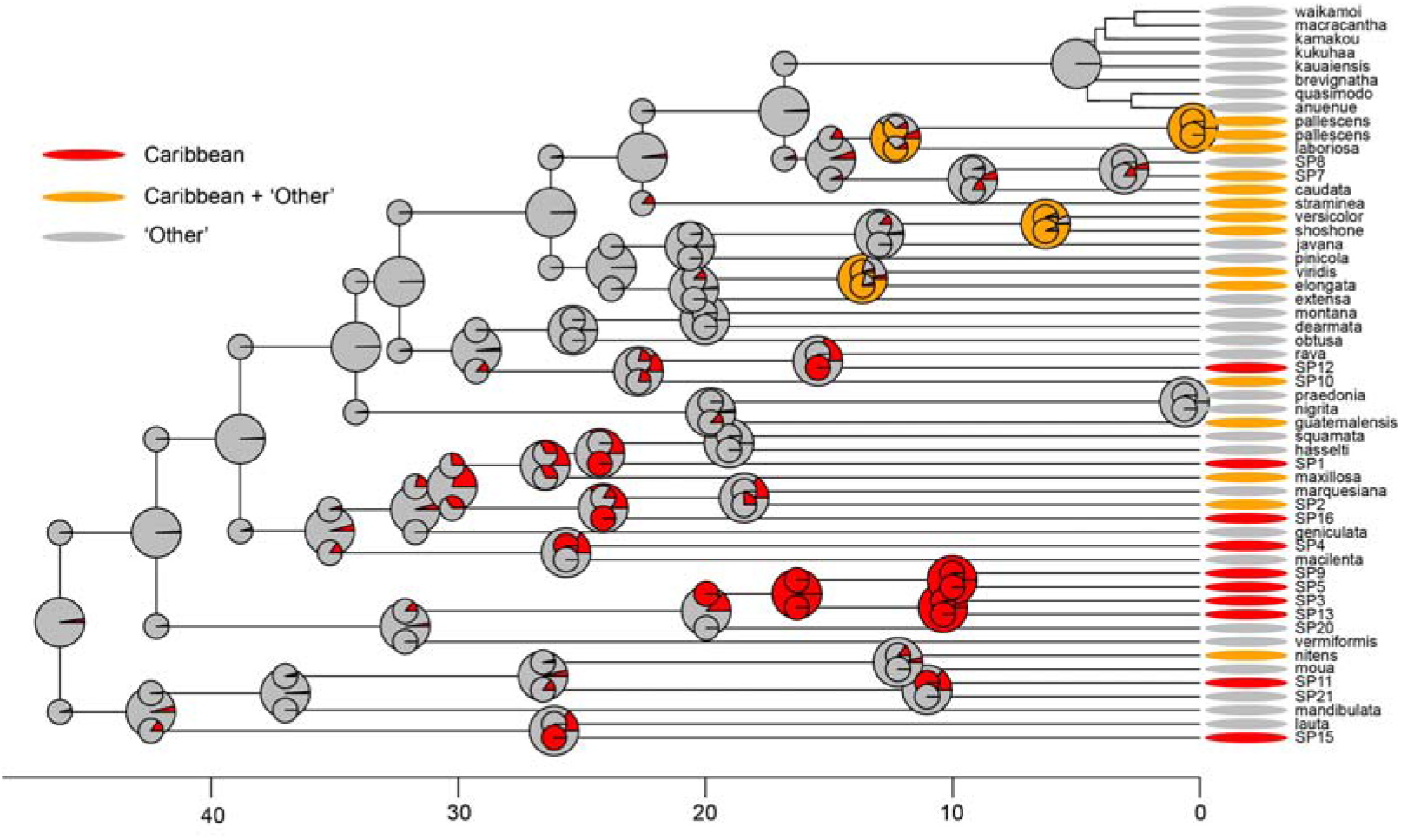
Ancestral area estimation of *Tetragnatha* with BioGeoBEARS. The biogeographical analysis using the most suitable model for our data (BAYAREALIKE + J, max_range_size = 2), reveals multiple origins of Caribbean taxa from the outside sources. A single subclade (SP 13, 3, 5, 9) has a well-supported Caribbean ancestral range. Evidently, multiple founder events and range expansion took place throughout *Tetragnatha* biogeographic history on the Caribbean.

We used 100 biogeographic stochastic mapping repeats of exact biogeographic histories (Animation A.1). This analysis estimated that on average 21.88 biogeographic events took place between the delimited areas. Of those, 11.72 were anagenetic events (all of which were range expansions), and 10.16 were cladogenetic events (Fig. 5). Of the latter, all were dispersals with founder event, and none were vicariant events. The results of narrow sympatry should not be considered in our case because of the experimental design with only two areas. Being extremely wide, the area classified as “Other” would produce a false positive score under narrow sympatry. According to BSM analyses 17.86 dispersal events must have taken place from outside sources to the Caribbean (9.51 range expansions, 8.35 founder events) while 4.02 dispersal events happened in the other direction (2.21 range expansions, 1.81 founder events).

**Figure 5.**
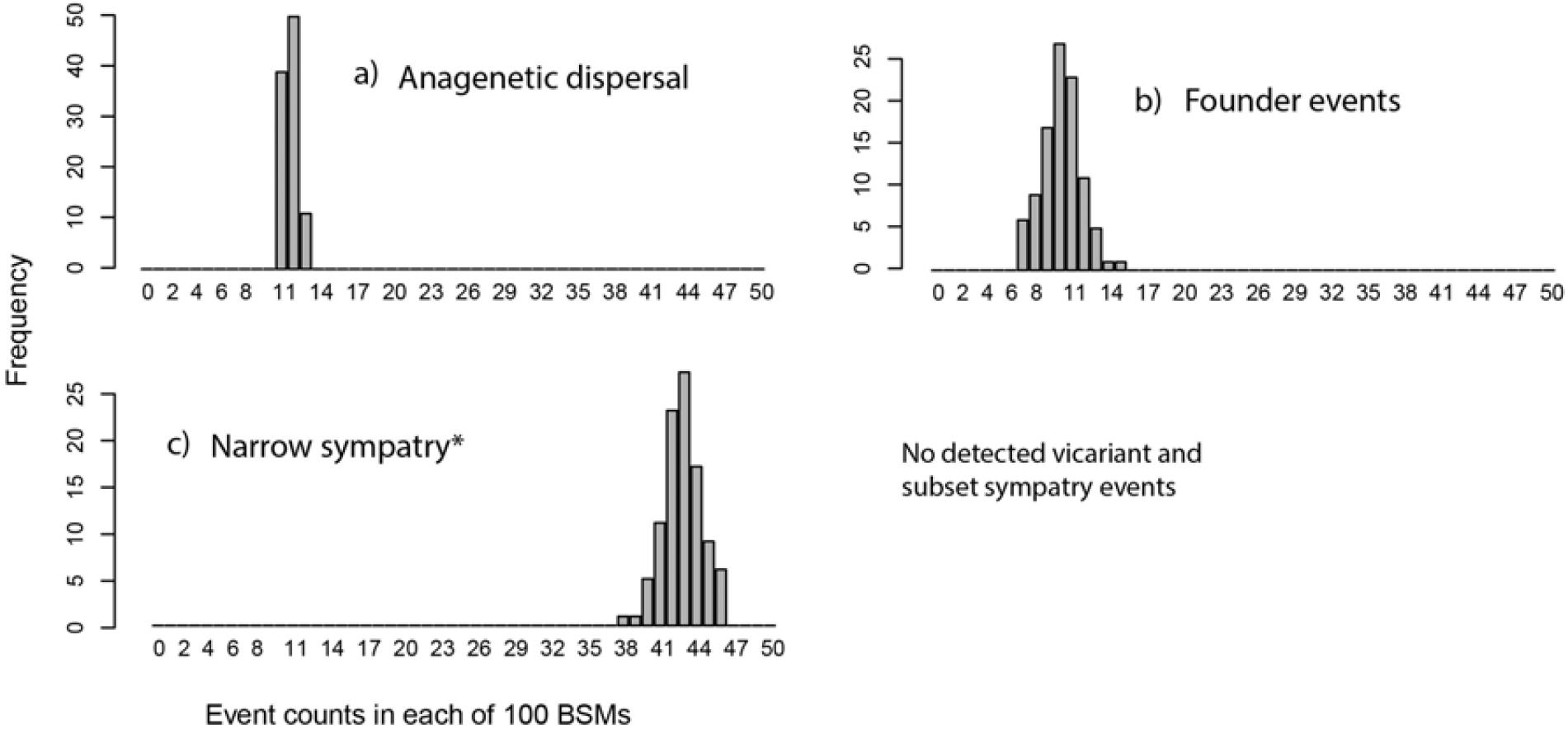
Histogram of event counts from Biogeographic Stochastic Mapping in BioGeoBEARS. Note that narrow sympatry should be interpret with caution due to experimental design with two areas. The area classified as “Other” is extremely wide and does produce a false positive score under narrow sympatry.

### 3.5 Tetragnatha SP2 haplotype network

Within the selected species *Tetragnatha* SP2, the network analysis recovered 21 haplotypes. The largest haplotype (VI) contained 16 individuals while eight haplotypes were represented by single individuals. Among haplotypes we counted 124 total mutations, with the longest link between two haplotypes having 20 mutations and the shortest a single one (Fig. 6).

**Figure 6.**
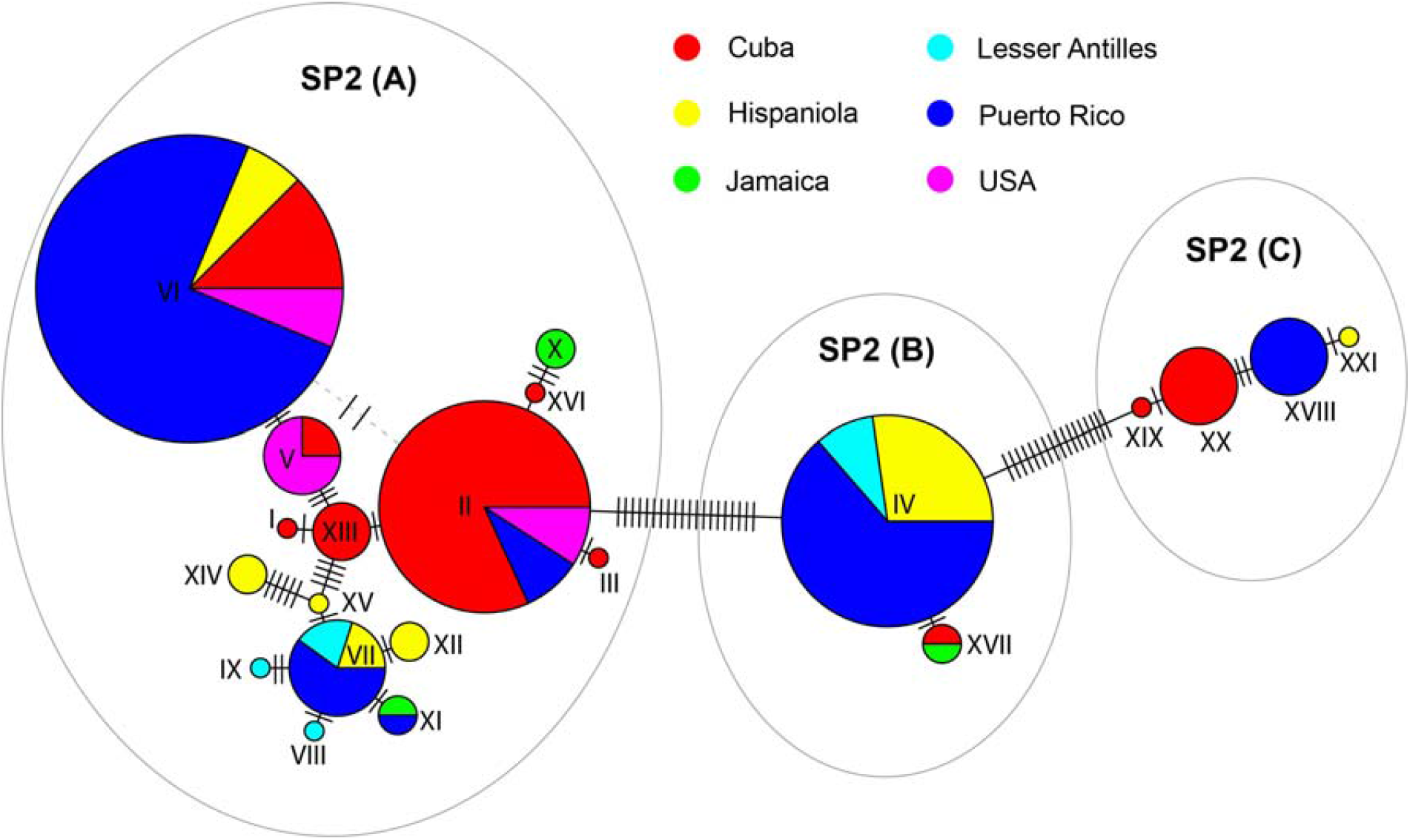
Haplotype network of *Tetragnatha* SP2 recovers three distinct haplotype groups. These groups (A, B, C) apparently do not intermix. Although our ABGD species delimitation lumped those three groups, the haplotype network strongly suggests that *Tetragnatha* SP2 contains cryptic species.

## 4 Discussion

We reconstruct multiple *Tetragnatha* phylogenies from over 300 individuals of this diverse genus. Our results support the monophyly of the four-jawed spiders but reject the monophyly of the Caribbean *Tetragnatha*. In the Caribbean, we find low levels of endemism yet high diversity within *Tetragnatha*, an unusual pattern considering other spider biogeographic research in the Caribbean (Agnarsson et al., 2018, 2016; Chamberland et al., 2018; Crews et al., 2010; Dziki et al., 2015; McHugh et al., 2014; Sánchez-Ruiz et al., 2015; Zhang and Maddison, 2012). The time calibrated phylogenetic reconstruction allows for an early overwater colonization of the Caribbean by *Tetragnatha* spiders. Moreover, the combination of chronogram and biogeographic history reconstruction refute the possibility of ancient vicariant origins of Caribbean *Tetragnatha* while favoring overwater dispersals rather than the use of GAARlandia land-bridge to reach Caribbean islands. Biogeographical stochastic mapping recovered multiple colonization events to the Caribbean and evidence of ‘reverse colonization’ from islands to continents, since mid-Eocene to late-Miocene. As we discuss below, our results, when compared with other lineages with known biogeographic histories in the Caribbean, suggest a unique dispersal history of *Tetragnatha*, combining excellent dispersal ability of the lineage as a whole with subsequent reduction or loss of that trait in individual clades through evolutionary history, as also seen e.g. in Hawaiian *Tetragnatha* (Gillespie et al., 2012, 1997).

### 4.1 Caribbean *Tetragnatha* are not monophyletic

Our phylogeny (Fig. 1) supports *Tetragnatha* monophyly, and provides a critical checkpoint for reliable estimation of MOTUs. However, combining our results with previously collected data is limiting, due to the lack of overlapping 28S data on GenBank. The comparison of this “local” phylogenetic reconstruction, involving exclusively Caribbean species, with the more taxon-rich, global phylogeny, highlights the importance of providing a global context to studies of Caribbean biogeography. While “local” relationships from Figure 1, based upon our sampling, are well supported and appear to tell a clear story, the picture changes drastically with the additional globally distributed species. Namely, the all-terminal phylogeny (Fig. 2) recovers a complex and mixed pattern in which single island Caribbean endemic species are phylogenetically scattered with other species, some of which are geographically distant. Overall, Caribbean *Tetragnatha* species do not form a clade. This Caribbean non-monophyly strongly hints at multiple colonization events of *Tetragnatha* to the archipelago. Moreover, the phylogeny did recover a single, small-scale, radiation of Caribbean *Tetragnatha* (the clade with SP3, 5, 9 and 13). This combined pattern suggests that at least some *Tetragnatha* species maintain relatively high levels of gene flow within and among the islands, as well as between the archipelago and the continents that serve as source populations. Others, however, seem to have secondarily lost this dispersal potential and form narrow range endemics. Similarly, researchers observed a lack of large monophyletic radiations of birds on the Caribbean (Ricklefs and Bermingham, 2008), that are otherwise well documented in bird lineages from more remote archipelagos such e.g. Darwin’s finches on the Galapagos (Grant and Grant, 2002) and Hawaiian honeycreepers (Lovette et al., 2002). On the other hand, the Caribbean island system does harbor its own exemplary, large, monophyletic radiation of *Anolis* lizards albeit a lineage with lower dispersal abilities than birds (Glor et al., 2004; Hass et al., 1993; Losos and Schluter, 2000). As it appears, the dispersal abilities of taxa must be ‘coordinated’ with isolation of an island or archipelago to provide the right conditions for an adaptive radiation (Ricklefs and Bermingham, 2008).

### 4.2 Biogeographic history of Caribbean *Tetragnatha*

Three hypothetical scenarios of Caribbean colonization are commonly reported. The first scenario, an ancient vicariant hypothesis, assuming the colonization of proto-Antilles as early as 70 MYA (Hedges, 2001), is refuted by our chronogram (Fig. 3). We estimate that *Tetragnatha* could have appeared on the Caribbean as early as 46 MYA, although more recent estimates are likelier (Figs. 3 and 4).

Therefore, the ancestral vicariance hypothesis receives no support by our data, because the scenario would predate our time estimates by roughly 30 million years, or more.

The second scenario assumes that non-marine organisms could have used the exposed land-bridge called GAARlandia to reach the Caribbean. GAARlandia, connecting the Greater Antilles with South America, possibly existed between 33 and 35 MYA (Ali, 2012; Iturralde-Vinent and MacPhee, 1999). While some research find support for colonization of the Caribbean via GAARlandia for diverse group of lineages e.g. fishes (Rícan et al., 2013), frogs (Alonso et al., 2012), mammals (Dávalos, 2004), invertebrates (Binford et al., 2008; Dziki et al., 2015; Matos-Maraví et al., 2014) and even plants (Fritsch, 2003; van Ee et al., 2008) but see (Nieto-Blázquez et al., 2017), we do not find evidence that would support the use of GAARlandia land bridge by *Tetragnatha*. Both, our chronogram and scattered phylogenetic pattern of Caribbean *Tetragnatha*, disagree with such scenario. Likewise, the pattern detected in our earlier study on *Cyrtognatha*, also refuted GAARlandia, but in that case the lineage was decisively too recent (Čandek et al. 2018 ‘in review’).

The third scenario involves overwater dispersal by terrestrial organisms to reach the Caribbean islands (Ricklefs and Bermingham, 2008). According to our time estimates (Fig. 3) and biogeographic history reconstruction (Fig. 4) we conclude that this scenario best explains our data. *Tetragnatha* (SP15) could have colonized the Caribbean as early as mid-Eocene, soon after the emergence of the first Caribbean islands between 40 - 49 MYA (Graham, 2003; Iturralde-Vinent and MacPhee, 1999; MacPhee and Grimaldi, 1996). Our phylogenetic (Fig. 2 and 3) and biogeographic (Fig. 4) history reconstructions suggest, that *Tetragnatha* repeatedly, and independently, colonized the Caribbean until mid-Miocene. Moreover, *Tetragnatha* biogeographic pattern within the context of the geological history of the Caribbean islands does not support a so called ‘progression rule’, a pattern where successive colonization of younger islands is correlated with cladogenesis (e.g. *Tetragnatha* on Hawaii; (Shaw and Gillespie, 2016)).

Evidence of multiple colonization events is strongly supported by our biogeographical stochastic mapping (BSM) analysis (Animation A. 1, Figure 5). Simulating biogeographic histories with BSM is a suitable approach to estimating the average number and directionality of biogeographic events in a studied area (Dupin et al., 2017). Within the scope of our analysis, we estimated that on average over eight founder events may have taken place to the Caribbean. Moreover, over nine times *Tetragnatha* species arrived to the Caribbean without subsequent speciation (anagenetic event of range expansion). Caribbean *Tetragnatha* also show evidence of reverse-colonization from islands to the mainland, likely with two range expansions and up to two founder events (Table A. 3).

Reconstructed biogeographic history of Caribbean *Tetragnatha*, is distinct from biogeographic patterns of *Tetragnatha* on other well studied archipelagos. Marquesas islands were probably colonized once (Gillespie, 2002), while Society islands have been colonized at least twice (Gillespie, 2002). On these remote Pacific archipelagos, *Tetragnatha* underwent monophyletic adaptive radiation although on a much smaller scale than *Tetragnatha* on the Hawaiian archipelago (Gillespie et al., 1997, 1994). *Tetragnatha* colonized Hawaii between two and four times (with two possible reverse-colonization events) and the subsequent adaptive radiation(s) resulted in at least 38 species (Casquet et al., 2015). On the other hand, the geographically less remote Mascarene islands in the Indian ocean, were colonized three times but did not undergo any adaptive radiation (Casquet et al., 2015). Considering the above examples, Caribbean *Tetragnatha* biogeographic pattern reveals (a) exceptionally high rates of colonization (and reverse-colonization), (b) relatively low levels of endemism, (c) generally more complex, phylogenetically scattered, species composition, and (d) relatively high species richness (only exceeded by Hawaii), compared to *Tetragnatha* from other archipelagos. It seems that the Caribbean offers a unique evolutionary arena, unlike any other.

### 4.3 *Tetragnatha* dispersal abilities

Our all-terminal phylogeny (Fig. 2), representing a global picture of *Tetragnatha*, recovers well-supported cases where Caribbean species group with geographically distant taxa, e.g. *Tetragnatha* SP15 from Hispaniola + *Tetragnatha lauta* from Asia; *T*. SP4 from Jamaica + *T. macilenta* from Oceania; *T*. SP12 from Hispaniola and Puerto Rico + *T. rava* from Oceania. Our taxon sampling may have omitted numerous intermediate species that may in fact group within these small but wide ranging clades. Nonetheless, these well-supported nodes may hold. Similar to our case, researchers found that *Tetragnatha* species amongst Pacific archipelagos (Marquesas, Society and Hawaiian) were geographically closer but phylogenetically more distant relatives than those between an archipelago and the mainland (Gillespie, 2002).

The extreme geographic distances (over 10,000 km) between pairs of closely related species in our phylogeny (Fig. 2), as well as often wide to cosmopolitan distributions of *Tetragnatha* species, together imply that *Tetragnatha* must contain numerous species with extraordinary dispersal abilities. On the other hand, the recurring pattern of single island endemism in *Tetragnatha* hints at evolutionary changes in this dispersal potential where certain species or clades within *Tetragnatha* secondarily have limited dispersal ability. The *Tetragnatha* as a genus thus exhibits high dispersal abilities and at the same time high intrinsic property to quickly adapt and diversify, having a “super-speciator” attributes (Linck et al., 2016; Pedersen et al., 2018).

To construct a more general picture of *Tetragnatha* dispersal ability, we compared biogeographic patterns of the Caribbean *Tetragnatha* with those of other Caribbean lineages of known biogeography and their estimated dispersal abilities in the theoretical context of the intermediate dispersal hypothesis model (IDM) (Agnarsson et al., 2014; Claramunt et al., 2012). In all cases, local (Caribbean) as well as global species richness of *Tetragnatha* is greater than in other genera with putatively poor dispersers such as: *Cyrtognatha* (Čandek et al. 2018 ‘in review’), *Deinopis* (Chamberland et al., 2018), *Micrathena* (McHugh et al., 2014), *Loxosceles* and *Sicarius* (Binford et al., 2008), *Spintharus* (Agnarsson et al., 2018; Dziki et al., 2015), *Selenops* (Crews and Gillespie, 2010). Moreover, attributes associated with putatively poor dispersers such as high levels of single island endemism and low numbers of colonization events are not reconstructed in the case of Caribbean *Tetragnatha*.

However, Caribbean *Tetragnatha* also show strikingly different patterns than spider genera of putatively excellent dispersal ability, such as *Nephila* (Kuntner and Agnarsson, 2011) and *Argiope* (Agnarsson et al., 2016). *Nephila* is distributed across the whole Caribbean archipelago but is represented by a single species, *N. clavipes*. Although several species of *Argiope* occupy the Caribbean, the most widespread is *A. argentata* (a recently discovered cryptic species *A. butchko* used to be *A. argentata* in part) (Agnarsson et al., 2016). This comparison, in addition to the previous comparison with putatively poor dispersers, reveals a much higher local (Caribbean) and global species richness in *Tetragnatha* than in *Nephila* or *Argiope*. Moreover, the biogeographic pattern with the combination of rare founder events, low number of widely distributed species, and extremely low endemism, all associated with an excellent dispersal ability, are not reconstructed in the Caribbean *Tetragnatha. Tetragnatha* is an excellent disperser, but appears to readily respond to natural selection upon colonizing island, which may render individual species to change their dispersal behavior and become locally endemic. While a direct test of the IDM would require consideration of three categories of dispersers, four-jawed spiders do not readily fit one of these three *a priori* definitions. Instead, they represent a more complex combination of attributes of a ‘dynamic disperser’.

### 4.4 Haplotype and cryptic species

Our analyses uncovered three distinct haplotype groups (A, B, C, Fig. 6) separated by 34 mutations in *Tetragnatha* SP2. Even though our ABGD delimitation analysis grouped them together, a closer examination reveals three potential cryptic species, requiring further morphological examination. Haplotypes from distinctively different groups (e.g. A and C) appear on the same island, but apparently do not intermix, a pattern that strongly indicates either an ongoing, or a completed speciation. In comparison, haplotype reconstruction of *Argiope argentata* and a cryptic species *A. butchko* (Agnarsson et al., 2016) had 23 mutations separating them while we detected 20 mutations between A and B groups (14 between B and C; 34 between A and C), further indicating the presence of cryptic species within *Tetragnatha* SP2.

A confirmation of cryptic species would call for an investigation of other widely distributed *Tetragnatha* species while further increasing the number of *Tetragnatha* species in the Caribbean to 28. Species richness of the Caribbean *Tetragnatha* is therefore much higher than that of another member of Tetragnathidae family, *Cyrtognatha* (~28 vs 11-14; Čandek et al. 2018 ‘in review’). Given that these genera are closely related, and co-distributed in parts of the region, the differences in species richness, as well as biogeographic patterns may, at least in part, be attributed to differences in their dispersal abilities.

## 5 Conclusions

Our research provides the first insight into Caribbean *Tetragnatha* evolutionary and biogeographic history. A combination of geographically scattered closely related species, single island endemics, cosmopolitan species, multiple dispersal events and high species richness of the Caribbean *Tetragnatha*, present a unique model for future comparative biogeographic research. Considering the theoretical framework of the IDM (Agnarsson et al., 2014; Claramunt et al., 2012), one cannot classify *Tetragnatha* as one of the three *a priori* definitions of dispersers (poor vs. intermediate vs. excellent). Instead, they represent a more complex combination of attributes of a ‘dynamic disperser’, one that readily undergoes evolutionary changes in dispersal propensity. Species richness, species distribution and disparate biogeographic pattern of *Tetragnatha* generally contrast with other arachnid lineages we have studied in the Caribbean. In line with the predictions of the IDM some highly dispersive *Tetragnatha* species fail to speciate on even remote islands. However, enough lineages have apparently secondarily lost dispersal ability, and thus formed short range (single island) endemics. Our findings demonstrate the very dynamic relationship between dispersal ability and diversity and how changes in dispersal propensity may link to endemism.

## Acknowledgements

We thank the entire CarBio team (http://www.islandbiogeography.org/participants.html) for collecting the material across the Caribbean. Moreover, we thank Lisa Chamberland and other members of the Agnarsson lab (http://www.theridiidae.com/lab-members.html) for the help with material sorting and molecular procedures.

## Data availability

All results generated in this study and protocols needed to replicate them are included in this published article, public repositories (GenBank) and supplementary material files.

